# Neural correlates in the time course of inferences: costs and benefits for less-skilled readers at the university level

**DOI:** 10.1101/2024.03.14.585085

**Authors:** Mabel Urrutia, Esteban J. Pino, María Troncoso-Seguel, Claudio Bustos, Pamela Guevara, Karina Torres-Ocampo, Sandra Mariángel, Yang Fu, Hipólito Marrero

## Abstract

Inferences are an indicator of a greater reading comprehension, as they imply a combination of implicit and explicit information that usually combines a textual representation with background knowledge of the reader. The aim of this study is to explore the costs and benefits of the time course of inferences in university students with reading comprehension difficulties at 3 stages during a narration. The method used was the event-related potential (ERP) technique in order to register the brain activity of 63 teaching program students while they read familiar, less-familiar and neutral stories. Results show a slow negativity potential component with greater negativity in words coming from familiar contexts when compared to less familiar and neutral ones in the first locus; an N400 component and a Post-N400 component in the second locus, reflecting greater negativity in familiar contexts when compared to less-familiar ones; and, lastly, through the use of a lexical decision task, FN400 and N400 components were found in the third locus, especially for pseudowords. These results are interpreted as a preferably bottom-up processing, which is characterized by lexical access difficulties in less-skilled readers.

## 1. Introduction

Inferences are essential in the comprehension of a text as they integrate global knowledge in the language model [1]. These must complete the limited information of the text with semantic and pragmatic knowledge available in the memory of the individual in order to make sense of the reading [2, 3]. In this research, we will study a type of bridging inference, causal coherence inferences, whereby the representation of general world, whether familiar or less-familiar, has an effect on reading comprehension [4], in particular for university students with reading comprehension difficulties.

An important characteristic of inferences is their double semantic representation: textual representation and background knowledge that is activated in parallel during inference processing. However, there is a singularity: one of the sources of information has an advantage over the other and determines the time course of reading comprehension [5]. Thus, familiarity of information coming from the knowledge of the reader and their capacity to integrate it into the text through inferences will make a difference in university-level readers [6, 7]. This dual processing of inferences implies the comprehension of implicit and explicit information [8, 9], whose level of activation varies according to reading skill. According to the results of the Programme for the International Assessment of Adult Competencies (PIAAC) [10], only 48.8% of young students worldwide make low-level inferences to comprehend texts. It is due to this that the study of this topic is of relevance to the field.

The main electrophysiological components that have been studied in inference generation are N400 and P600. The N400 component is a negativity that appears in the temporoparietal region between 200 and 600 ms when facing the probability of appearance of an stimulus. On the other hand, the P600 component is a positivity that appears in the parietal region at around 500 ms when facing semantic review and repair processes; a re-analysis that is both syntactic and semantic due to an error in the interpretation of the phrase being read [11]. Some current studies have described another component called post-N400, which relates to the frontal-posterior regions and that relates to a response arising from a semantic violation surrounding a very specific context [12]. However, there is yet no clarity as to the behavior of such component.

Electrophysiological measures have allowed determining the temporal stages of cognitive processing in inference solving. Thus, effects have been reported in the N400 component when facing a semantic problem induced by discourse not fitting with the broader context of the story [13]. Additionally, the N400 component has been found when presented with the manipulation of incoherent inferences in a social context [14, 15]. Tabullo et al. [16] found an N400 component associated to unexpected endings coming from highly restrictive contexts during phrase reading.

On the other hand, the P600 component has been associated with context updating processes during inference processing [14]. Burkhardt [17] also found a higher positivity, associated to the P600 components, in probable and inducible contexts in an experimental manipulation related to phrase reading. Kuperberg et al. [18] found a P600 component in anomalous critical words that violated the restriction of the context, whether it was in low context or in a highly restrictive context, when compared to plausible critical words.

On the other hand, some experimental evidence with ERP show that when the context is tightly related to the target word, the prediction of said word was high and, as a result, the amplitude of the N400 was lower [19]. Kuperberg et al. [20] studied causal relationships in different scenarios. Their results show a higher amplitude of the N400 component in causally unrelated scenarios when compared to highly and intermediately causally related scenarios, regardless if they appeared before or after the final position of the sentence. This shows the effect of contextual information on inference making. Ito et al. [21] showed an attenuation of the N400 component for highly predictable as well as semantically related words when compared to unrelated words. Thus, the amplitude of N400 is modulated by the reader’s difficulty to integrate lexical information in a predictable context.

In spite of that, a type of N400 with a topographically left-frontal distribution has been discovered: the FN400 component, directly linked to familiarity in semantic language processing [22]. This is a neural correlate of familiarity that has been found on several recent investigations [23, 24, 25, 26]. In Mecklinger and Bader’s [27] meta-analysis, there is a discussion on a series of experiments that show this component as a quick evaluation of the fluency of ongoing processing, in relation to previous events or current expectations, which are related to the mechanism of relative familiarity. During an episodic recognition test, Leynes and Upadhyay [25] found the FN400 component in pre-experiment familiar names, while pre-experimentally unfamiliar names evoked no old/new difference during the 300 to 500 ms interval. Bade et al. [26] found the FN400 component in a recognition test where the effect of a previously named word or a new word was manipulated through a semantic judgment. The results support the notion that the FN400 takes place when fluency is attributed to familiarity during a recognition decision test.

One of the main inspirations of this research was that of Steele et al. [7], who manipulated the familiarity variable in 3 contexts: familiar, less-familiar or neutral. After reading the phrases, a lexical decision task was done using words the participants may have inferred from the context or words that were unrelated to the scenarios. The results showed a reduction of the N400 component for words coming from unfamiliar contexts, as well as late positive activity. This was linked to the P600 for words coming from a familiar context when compared to a neutral context. The authors interpret such results as a marker of causal coherence inferences. In this research, we follow their experimental paradigm, with certain modifications to the SOA for the sake of contrast. Likewise, this research follows the time course of causal inferences starting from the context, through the critical phrase and up to the target; in three different moments of discourse.

Inferences that establish causal coherence are processed rapidly and automatically according to some authors [28]. The time course of such inferences is accelerated by context restrictions and, as a result, they are more likely to be generated [29]. The manipulation of inferential context familiarity will evoke fewer alternatives for a representation of the event, therefore leading to a faster generation of inferences [30]. As the inferences in question are bridging or forward inferences —since the target word is related to the context of the story—, such inferences adjust better to McKoon and Ratcliff’s [31, 32] minimalist hypothesis. The authors’ theory posits that inferences are produced online only if they are supported by information coming from experience according to the situation model on general background knowledge.

### The study

The traditional position is that a less-skilled reader considers reading as a decoding task, while a skilled reader understands it as a meaning-building task [33]. Less-skilled readers have difficulties in particular while comprehending the vocabulary of difficult and meaningless words. Additionally, there are external consistency difficulties in the detection of information violating prior knowledge. On the other hand, they face difficulties in the detection of errors affecting the internal coherence of the text, especially in expository texts. In this regard, it is particularly difficult for less-skilled readers to access potential background information in the representation of discourse in order to build a coherent representation of the text during the course of reading [34]. In the case of Chilean university students, only 10% of them rank in the highest levels of reading comprehension [35]. The main difficulties they face is with implicit questions, such as linking information, making inferences, integrating information or transferring information coming from general knowledge to finally determine the global coherence of the text [36].

The general aim of this research is to explore the costs and benefits of inference time course in university students with reading comprehension difficulties. The hypothesis of this study is that context in stories will influence the students from the start of the narration, eliciting differences between familiar and less-familiar contexts in the N400 component. On the other hand, due to reading difficulties of the participants, lexicality of the words and pseudowords will interact with the familiar and less-familiar context in the generation of inferences, with a possible FN400 or N400 effect.

## 2. Materials and Methods

### 2.1. Experiment Design

To test the hypothesis of the study, a within-subject 3 (high familiarity/low familiarity/neutral context) x 2 (word/pseudoword lexicality) factorial design was defined. Subject groups were randomly assigned and described based on each level of the variables. The independent variables were the familiar (F), less-familiar (LF), and neutral (N) context, and the word or pseudoword presented as target. The dependent variable is the amplitude of the components in ERP responses.

### 2.2. Participants

Sixty-three adults (56 women) ranging in age from 19 to 30 years (M=21.3, SD= 2.31) volunteered for the experiment. The participants were students from the Faculty of Education at the Universidad de Concepción (Chile) who volunteered to participate in the experiment. All volunteers were undergraduate university students who were pursuing a teaching program at the Universidad de Concepción (Concepción Campus).

The students were enrolled in an elective course for improving reading comprehension, which offered academic credits. Two diagnostic tests on reading comprehension revealed that 66% of the males and 68% of the females were below the reference score of the Inter-American Reading Test [37], which measures vocabulary, cloze, reading of short texts. Also, the students obtained a low percentage of reading comprehension and were ranked between the 78th and 79th percentile of the reference score of the Lectum test [38], which measures long texts at a discursive level. Lectum is a diagnostic test used at the students’ university.

For the estimation of the minimum required sample size, the following parameters were considered for a one-way analysis of variance test: a) Effect size (f)=0.25, b) Statistical power (1-β)=0.95; c) Significance level (α)=0.05; d) Number of measurements=5. According to these variables, a minimum of 31 individuals per group was needed, as calculated by the G*Power program version 3.1.7. In order to address university dropout effect, which is common at this educational stage, we have considered a sample size almost double that which is required. From the original 63 participants, data from five participants were removed: one participant due to technical problems during experiment execution, four for having a low percentage of correctly recorded trials. Data from 58 participants were used for the analysis (51 women).

Additionally, the following exclusion criteria were considered: (a) Having a history of uncorrected sensory impairments (e.g., vision), (b) having a history of neurological disorders (e.g., epilepsy, migraines, etc.), (c) having a history of substance abuse, (d) having a history of learning disorders (e.g., dyscalculia), (e) having a history of language disorders (e.g., dyslexia, aphasia, etc.). All participants were right-handed with normal or corrected-to-normal vision. They received monetary compensation for transportation expenses. All participants received information on the experimental procedure and gave their written informed consent before participating. Data recording began on August 1, 2022 and ended on September 9, 2022.

### 2.3. Stimuli

The material for the inference task was adapted from the study of Sundermeier et al. [7]. A normative study was carried out to assess context familiarity of texts. Participants were asked to determine the degree of familiarity on a scale of 1 to 7, where 1 was LF and 7 was highly F. The results showed statistically significant differences between high and low familiarity texts t(59)=4.421, p= 0.0001. In addition, a normative study cloze probability was conducted to determine the probability of the target word appearing in the experimental texts. Participants were asked to answer on a scale of 1 to 7 how likely it was that the target word would appear in the text, where 1 was very unlikely and 7 was highly likely. The results showed that there were no significant differences between the variables high familiarity and low familiarity. Therefore, in both contexts the target word could be inferred t(59)= 1.955, p= 0.057. All words and the sentences endings were controlled for their length between F and LF sentences. Thus, in the context sentence, there were no significant differences in length t(59)= -1.187, p= 0.240; in the critical sentence t(59)= 0.362, p= 0.240 and also in word target t(59)= 0.897, p=0.374. In the same vein, the frequency of use of the words of the sentences was controlled according to Chilean lexical frequency [39]: final word of context sentence t(59)=0.951, p=0.346; final word of critical sentence t(59)=1.528, p=0.132; target word, t(59)=1.283, p=0.204. Finally, we controlled for the variable imaginability, finding no significant differences in any of the final words in the sentences: the context sentence, t(31)=1.447, p=0.158; critical sentence, t(31)=0.510, p=0.614; target word, t(31)=-1.690, p=0.103.

Experiment Builder software (version 2.3.1; SR Research, 2020) was used for stimulus presentation. The experiment consisted of 93 stimuli, made up by two sentences and a target word. The first 3 stimuli were part of the training section and were not used for the analysis of the results. The sentences were strategically structured to end with a critical word, where the experimental manipulation related to familiarity was reflected (context: F, LF, and N). The target word was presented in the center of the screen, and participants were tasked with a lexical decision, determining whether the stimulus was a real word or a pseudoword through a YES/NO response. In 30 out of the 93 sentences, a comprehension verification question was included to assess text comprehension and ensure participants’ attention to the task (see **Table 1**). Stimuli were presented randomly within 3 blocks of 30 texts.

**Table 1.** Example stimuli presentation.

| Context | Sentence No. | Sentences | Presentation |
| --- | --- | --- | --- |
| Fixation point |  | + | 1000 ms |
| **Familiar** | 1 (context) | Para preparar el examen, el estudiante abrió su libro.<br>(*To study for the exam, the student opened his book.*) | Self-administered<br>(up to 3500 ms) |
|  | 2 (critical) | Después de tres horas, estaba seguro de que conocía muy bien la materia.<br>(*After three hours, he was sure that he knew the material well.*) | Self-administered<br>(up to 3500 ms) |
| **Less-familiar** | 1 (context) | El agente de la PDI recogió la fotografía del ladrón.<br>(*The police officer pick up the thief's photograph*)<br>(*After three hours, he was sure that he knew the material well.*) | Self-administered<br>(up to 3500 ms) |
|  | 2 (critical) | Después de unos minutos, se sintió seguro de que podría identificar al hombre en la calle.<br>(*After a few minutes, he felt confident that he could identify the man on the street*) | Self-administered<br>(up to 3500 ms) |
| **Neutral** | 1 (context) | Los niños llegaron a su primer día de escuela.<br>(*The children arrived at their first day of school*) | Self-administered<br>(up to 3500 ms) |
|  | 2 (critical) | Estaban emocionados de conocer a su nuevo profesor, hacer nuevos amigos y jugar en el recreo.<br>(*They were excited to meet their new teacher, make new friends, and play at recess*) | Self-administered<br>(up to 3500 ms) |
| **Target word** | Task | **ESTUDIO**<br>(STUDY) | Self-administered<br>Self-administered |
| **Yes/No question** |  | Is it a word? Yes/No answer through button box<br>(word/pseudoword evaluation) |  |
| **Post sentence blank screen** |  |  | 1500 ms |

\* Tables may have a footer.

### 2.4. EEG Data Recording

During the experiment, EEG data were collected using a sixty-four active electrode actiCAP elastic cap (FP1, FP2, AF7, AF3, AFz, AF4, AF8, F7, F5, F3, F1, Fz, F2, F4, F6, F8, FT9, FT7, FC5, FC3, FC1, FC2, FC4, FC6, FT8, FT10, T7, C5, C3, C1, Cz, C2, C4, C6, T8, TP9, TP7, CP5, CP3, CP1, CPz, CP2, CP4, CP6, TP8, TP10, P7, P5, P3, P1, Pz, P2, P4, P6, P8, PO7, PO3, POz, PO4, PO8, O1, Oz, O2, Iz), positioned according to the international 10/10 system. The reference electrode is located in the central area of the cap (FCz), and the reference electrode on the forehead (BrainProducts).

To set up the experiment, the electrodes were first connected to the actiCAP ControlBox to asses contact impedance. This set-up was performed for each participant, to ensure that all 64 channels’ impedance was lower than 25 KΩ before running the experiment.

The recording of the data for each participant was performed by connecting the electrodes to a standard BrainAmp amplifier, which was isolated from the power supply line by means of Brain Products Power Pack batteries. Each amplifier was connected, via fiber optic, to a USB-BUA128 adapter that transferred the signals in real time via USB port and cable to a laptop running BrainVision Recorder capture software. Data was recorded using a sampling rate of 500 Hz.

The stimuli were displayed on a DELL computer, model G7 7790, featuring a 17.3-inch screen (1920x1080 pixels) and a refresh rate of 60 Hz. We used Experiment Builder software from SR Research for configuring and presenting the stimulus.

A response button box was used to record participants’ responses. One button was used as ‘enter’ to finish self-administered steps. A ‘yes’ and a ‘no’ button were used to respond to the lexical decision (word/pseudoword) and to the attention questions.

Synchronization signals were sent from Brainvision Experiment Builder to Brainvision recorder to mark the display of the target on screen and at the participants’ response to the lexical decision.

### 2.5. Procedure and Experimental Paradigm

In the experimental session the appropriate size of the EEG cap was selected to fit each volunteer’s head. For the reading task, participants were seated in a height-adjustable chair, positioned in front of the stimulus presentation computer, which was placed at a distance of 70 cm from the participant.

General instructions for the experiment were displayed on the screen. The experiment was divided into three blocks, each containing 30 stimuli, with breaks provided between each block to allow participants to rest. The experimental session lasted on average 40 minutes per participant. At the start, three additional stimuli served as practice session. Stimuli were presented randomly within each block, with an equal distribution of F, LF, and N context (1/3 each). All experimental contexts were counterbalanced across participants. A lexical decision task was used to determine if the target words were words or pseudowords. Half of the presented stimuli were words and half were pseudowords. An example stimulus is shown in Figure 1.

**Figure 1.**
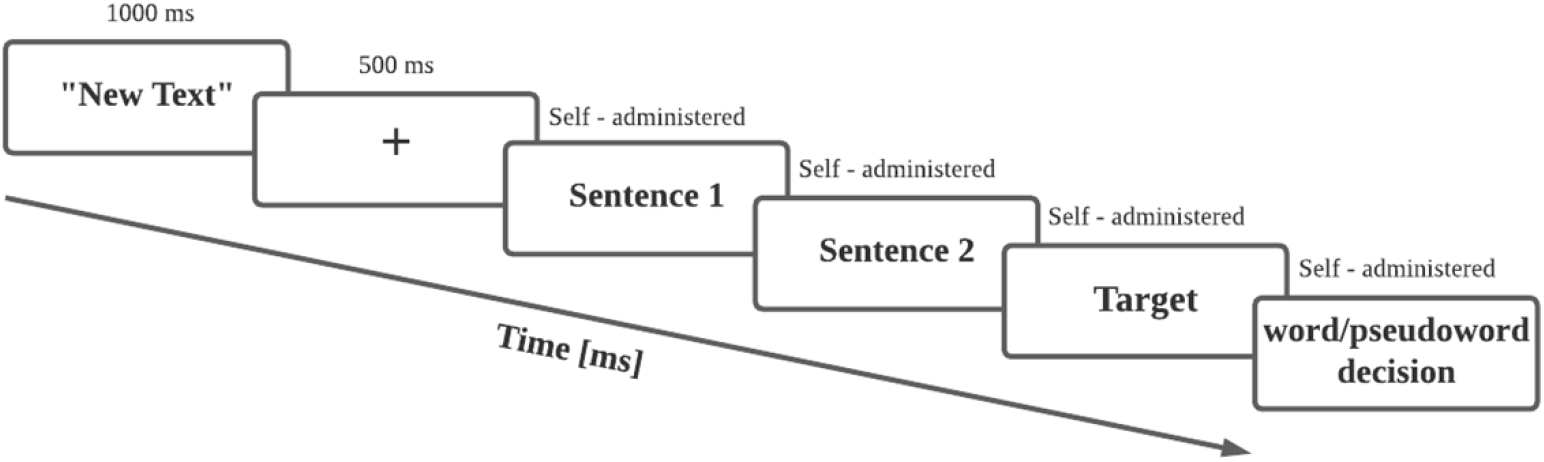
Experimental sequence diagram for each stimulus.

### 2.6. Data Analysis

A quantitative analysis was conducted, including descriptive statistics (measures of central tendency and dispersion), as well as inferential statistics using a linear mixed-effects model on the averaged data with random intercept for subjects. Statistical power and effect size are reported for the results obtained.

The EEG data analysis was performed using Brain Vision Analyzer software to analyze the brainwaves for all stimuli. All analyses were performed with R software (version 4.3.) with the ‘lme4’ package for linear mixed-effects modeling [40].

For ERP analyses [Y], a mass univariate analysis was carried out to select the specific time windows. A point-by-point t – test analysis was used for the whole epoch, which was then visually inspected to define the time window. For the analysis of locus (1 and 2) we only included the context as fixed effect. However, for the analysis of locus 3 (target word), we included the interaction between context (F, N, LF) and lexicality factors (word, pseudoword). For random effects, an intercept-only model is used, which represents the subject effect.

We used the anova output of lmer (Type III with Satterthwaite’s method considering a significance level of 5%) and R emmeans package to analyze contrasts. Contrasts were made with the emmeans packages using the pairwise method and p values were adjusted with the Tukey method.

For reaction times in the behavioral analyses of the lexical decision task, the same mix-model used in locus 3 analyses was applied, with the difference that the trial is added as random effect. Additionally, we used the anova output of lmer and explored contrasts. Similarly, for the analysis of task accuracy, by used a generalized mixed-model with binomial family considering correct and wrong answers in the model. For random effects, an intercept-only model was used, which represents the subject effect. We presented the summary and anova output results of the model.

#### 2.6.1. EEG Data Pre-processing

EEG data were processed using Brainvision Analyzer v2.3. The pre-processing pipeline included a visual inspection of EEG channels [41, 42, 43, 44]. In case of channel rejection, the channel was interpolated from neighboring electrodes. Then, all channels were re-referenced to the mean, and all channels were subsequently filtered from 0.1Hz to 30Hz with an 8th order FIR filter, followed by a notch filter. Then, a visual inspection to detect artifacts using a semiautomatic method with a threshold of 150uV and a gradient of 50uV/ms was done. Also, ocular artifact correction using ICA from Brainvision Analyzer was used, with FP2 (vertical) and F8 (horizontal) references. EEG data were segmented into epochs of interest: -200 to 1000 ms time-locked to the target word onset. The individual mean was calculated grouping by context (F, N, LF) and by lexicality (word, pseudoword), without considering the segments that were marked as faulty during visual inspection. Then, a baseline correction between -200 to 0 ms was applied. Finally, epochs were averaged to obtain the grand average for critical word (locus 1 and 2) and target stimulus (locus 3). These were later grouped by context and by lexicality for visual exploration of the different components.

## 3. Results

In this section, we show the results of the analysis of each type of the studied locus: locus 1, locus 2 and locus 3 in three different segments of the text.

### 3.1. Locus 1: slow negativity potential

At the critical word 1, we found a slow negativity potential for the ROI comprising electrodes FT8 – FT10 – T8 – TP8 – TP10 of the right hemisphere, as shown in Figure 2 and Figure 3. Both the right frontal dorsolateral topographical distribution and the late potential in the window where the effect can be seen are evidence of the component.

**Figure 2.**
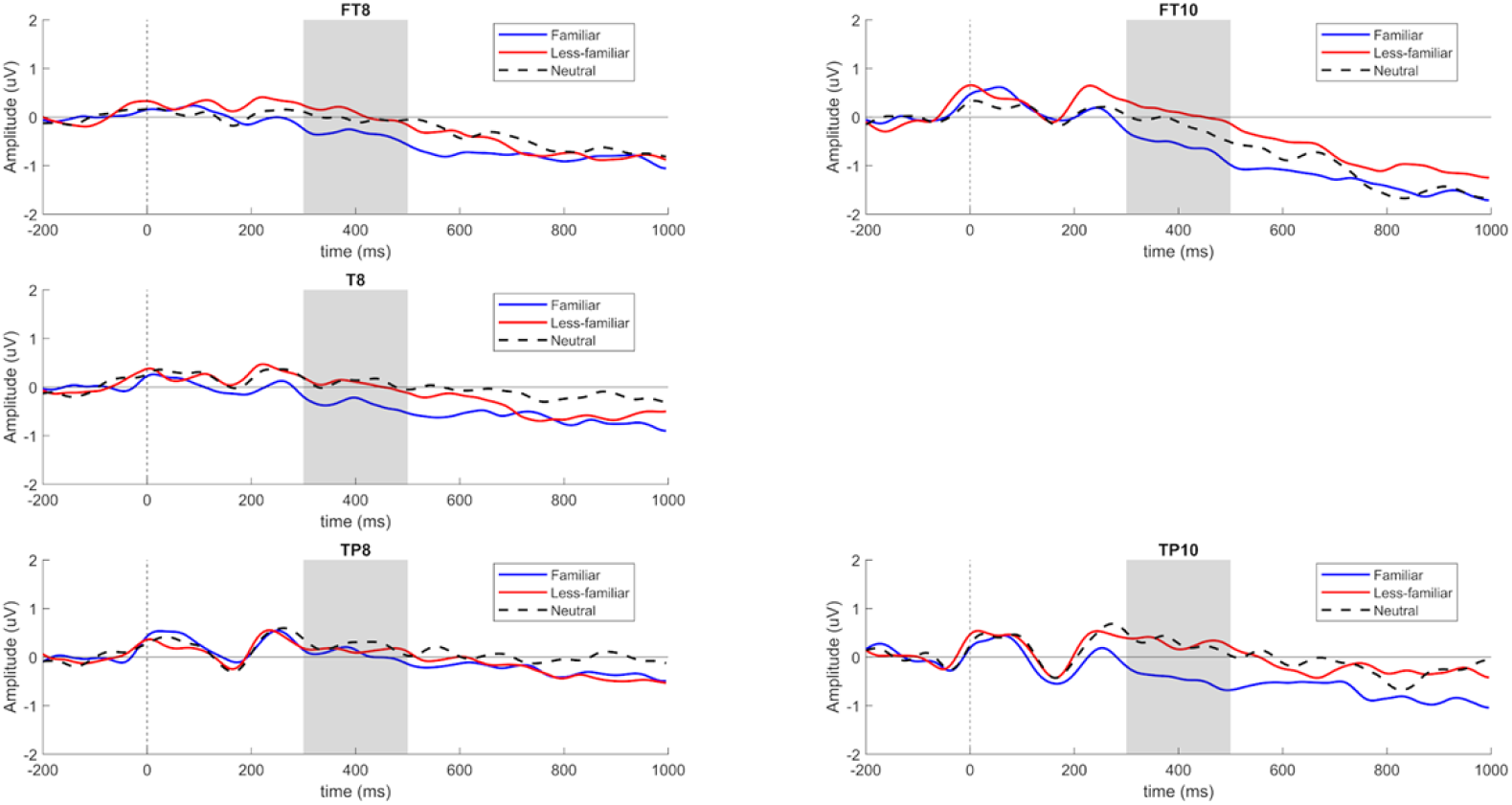
**Slow Negativity Potential component** of Locus 1 for electrodes FT8 – FT10 – T8 – TP8 – TP10 sorted by F, N and LF conditions.

**Figure 3.**
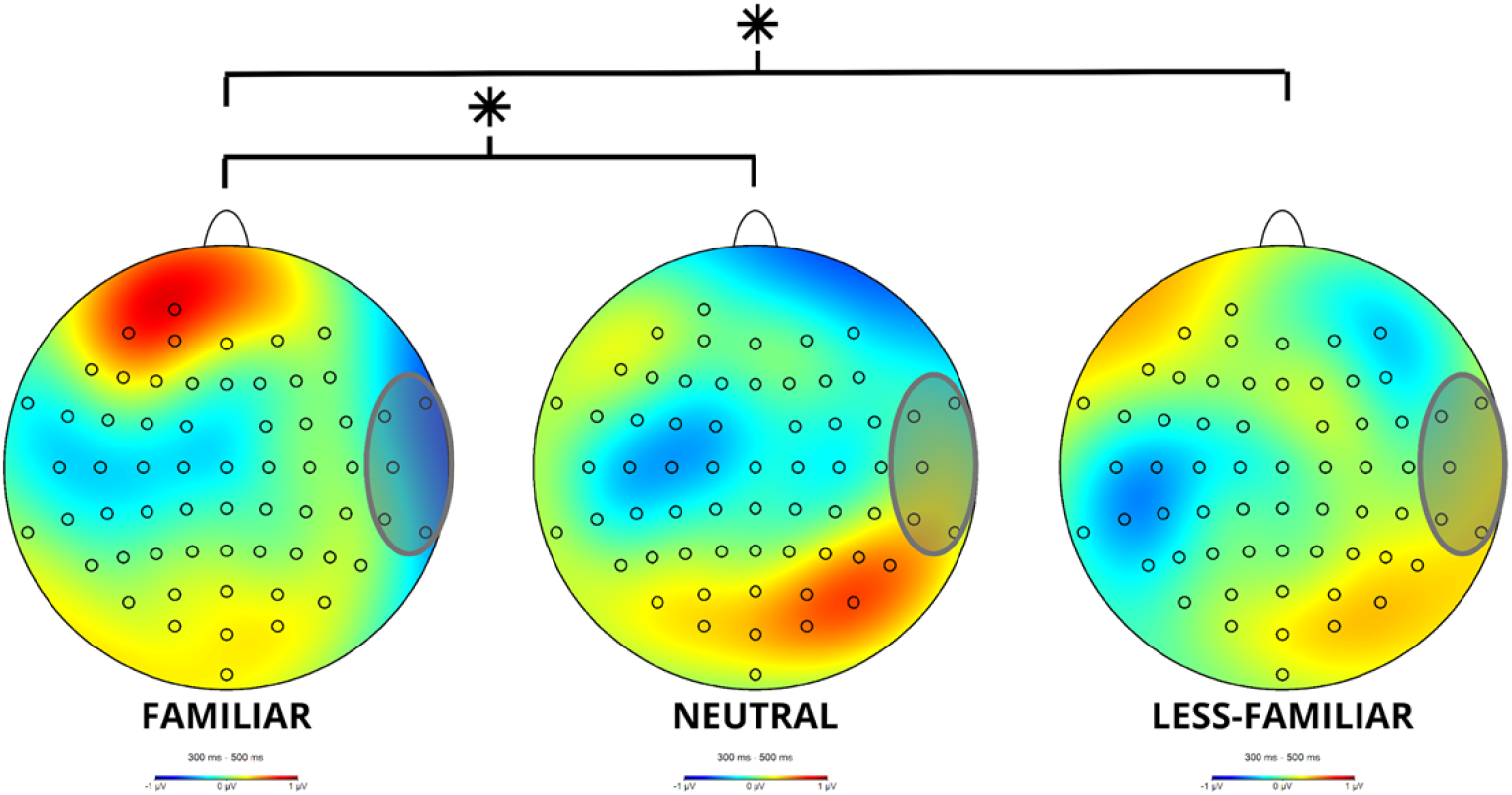
Topographical map of the **Slow Negativity Potential component** in Locus 1 in the studied 300-500 ms time window for the F, N and LF conditions.

The analysis of the 300-500 ms time window shows significant differences F(2,114)=4.368, p=0.015, for the analyzed conditions (F, N, LF).

The contrasts show a significant difference in the comparison between F and LF, t(114)=2.707, p=0.021 with a more negative amplitude for F than for LF. Additionally, a significant difference was found between F and N, t(114)=2.381, p=0.049, where the F context was more negative than N.

As shown in Figure 2, the familiar experimental conditions show higher negativity when compared to the less-familiar and neutral conditions.

### 3.2. Locus 2: N400 and Post N400

#### N400

For critical word 2, an N400 component was detected in the left central parietal region, involving electrodes C1, C3, C5, CP1, CP3 and CP5, as shown in Figure 4. Figure 5 shows the topographical map for each of the conditions.

**Figure 4.**
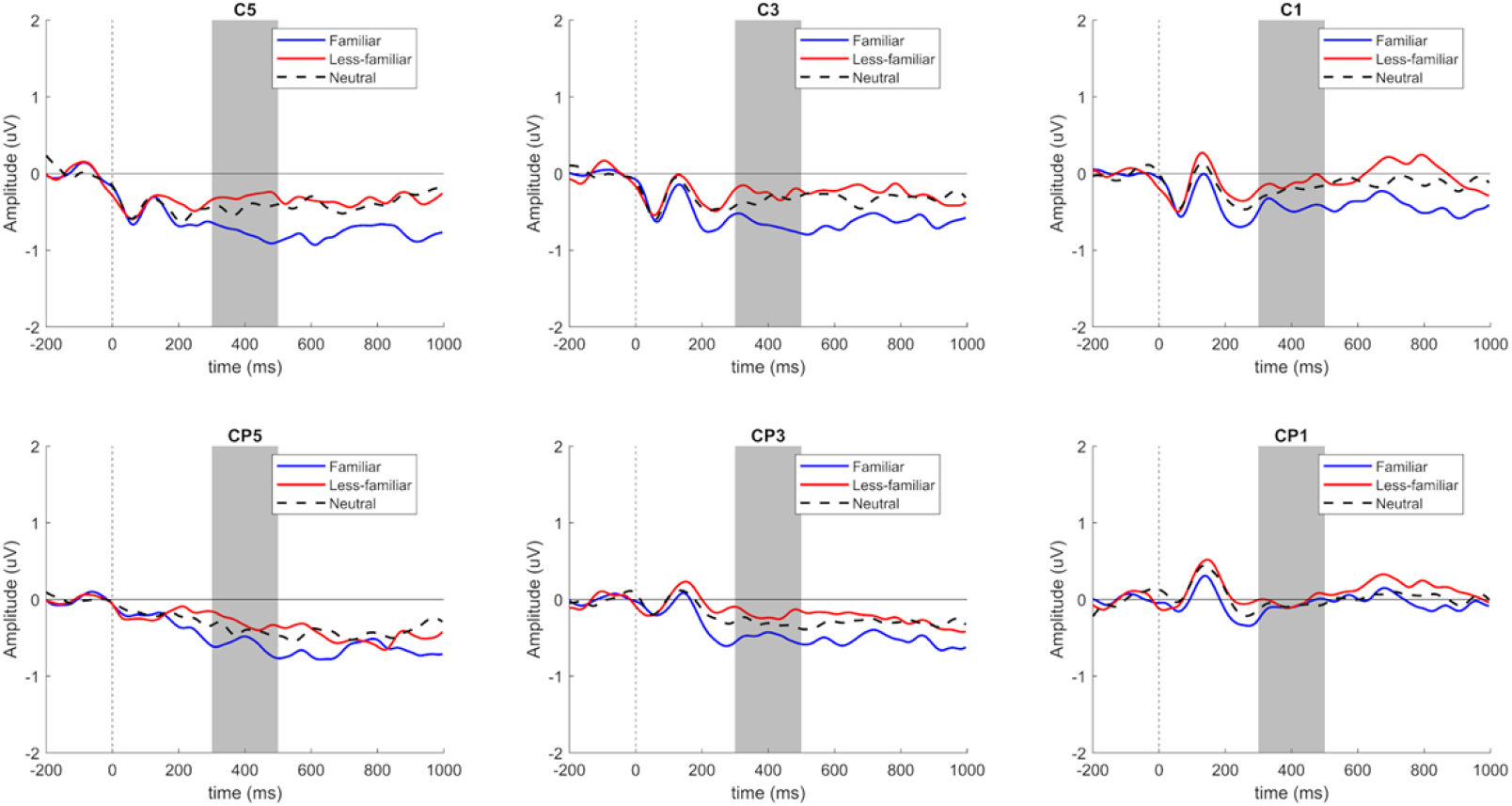
**N400** component of Locus 2 for electrodes C1 – C3 – C5 – CP1 – CP3 – CP5, sorted by F, N and LF conditions.

**Figure 5.**
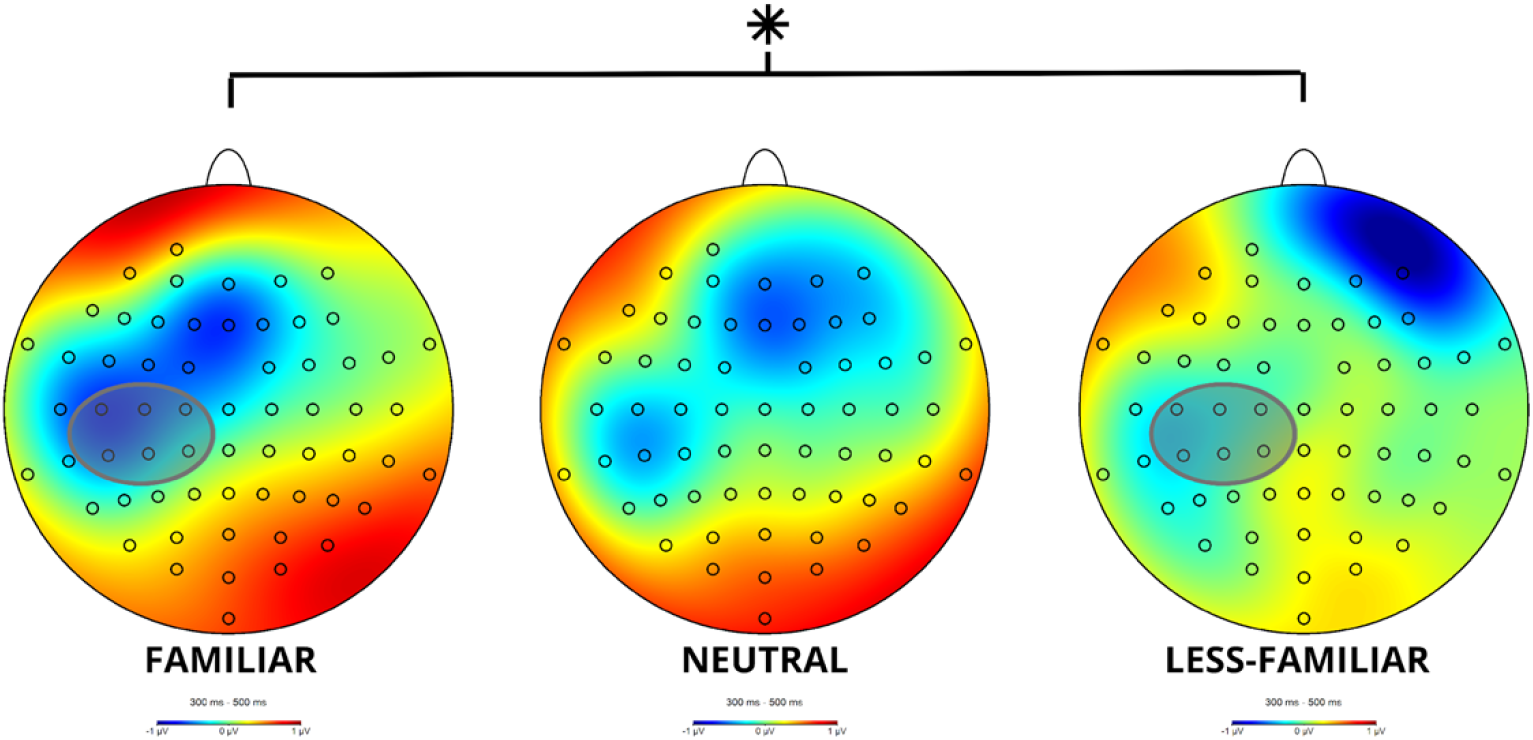
Topographical map of N400 component for Locus 2 in the 300-500 ms time window for the F, N and LF conditions.

Based on the ANOVA analyses, we found a significant difference F(2,114)=3.47, p=0.035 in the analyzed contexts. The contrasts show a significant difference between the conditions F and LF, t(114)=-2,59, p=0.029.

As shown in Figure 4, there is a higher cognitive cost in the familiar context when compared to the less-familiar context. Likewise, the topographical map shows this negativity with a greater left distribution in the familiar context when compared to the other experimental conditions (Figure 5).

#### Post N400

In locus 2 at the 600-1000 ms time window a post-N400 component was found in the ROI, involving electrodes FC3, FC1, FC2, FC4, C3, C1, Cz, C2, C4, CP3, CP1, CPz, CP2 andCP4. The analysis of this time window shows a significant difference, F(2,114)=4.50, p=0.013, for the analyzed conditions.

Contrasts reveal a significant difference between conditions F an LF, t(114)=-2.94, p=0.011.

### 3.3. Locus 3

#### 3.1.1. Behavioral results: Reaction Time and Accuracy

##### Reaction Time

At a behavioral level, the interaction of the variables context and lexicality was not significant, F(2, 4801.3)=1.02, p=0.36. However, as a main effect, lexicality is statistically significant, F(1, 4804.8)=645.72, p < 0.001, i.e. participants took longer to respond to pseudowords than to words, as expected. The context main effect was not significant, F(2,4801.3)=2.47, p=0.084. As there is a lexicality main effect, we can infer that both variables, word and pseudoword, behave differently. As a result of this, there is no significant context main effect.

Since the interaction was not significant, we explored the results separately by word and pseudoword in order to look at the behavior of each experimental condition. This was done with the same aforementioned method (lmer).

A significant difference was found in relation to the context of the word, F(2,2360)=5,23, p=0,005. Contrasts reveal a significant difference between F and N, t(2358)=3.076, p=0.006, in which students took longer to respond to F than to N. Likewise, there was a significant difference between LF and N, t(2359)=2.420, p=0.0412, in which participants took less time to respond to LF words than to N. Results are displayed in Figure 8.

**Figure 6.**
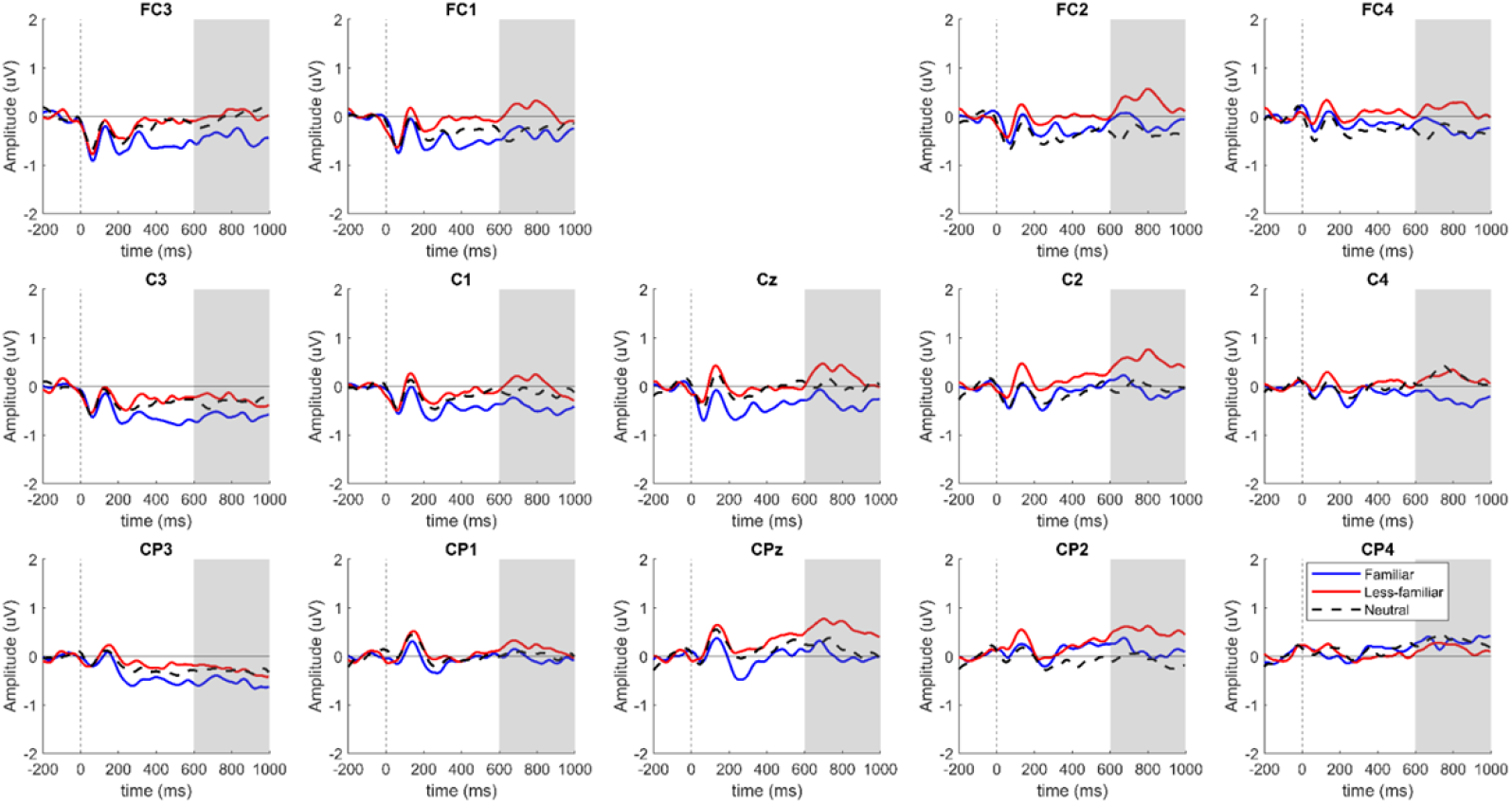
Locus 2 **Post N400** Component for electrodes FC3, FC1, FC2, FC4, C3, C1, Cz, C2, C4, CP3, CP1, CPz, CP2, CP4 sorted by F, N and LF conditions.

**Figure 7.**
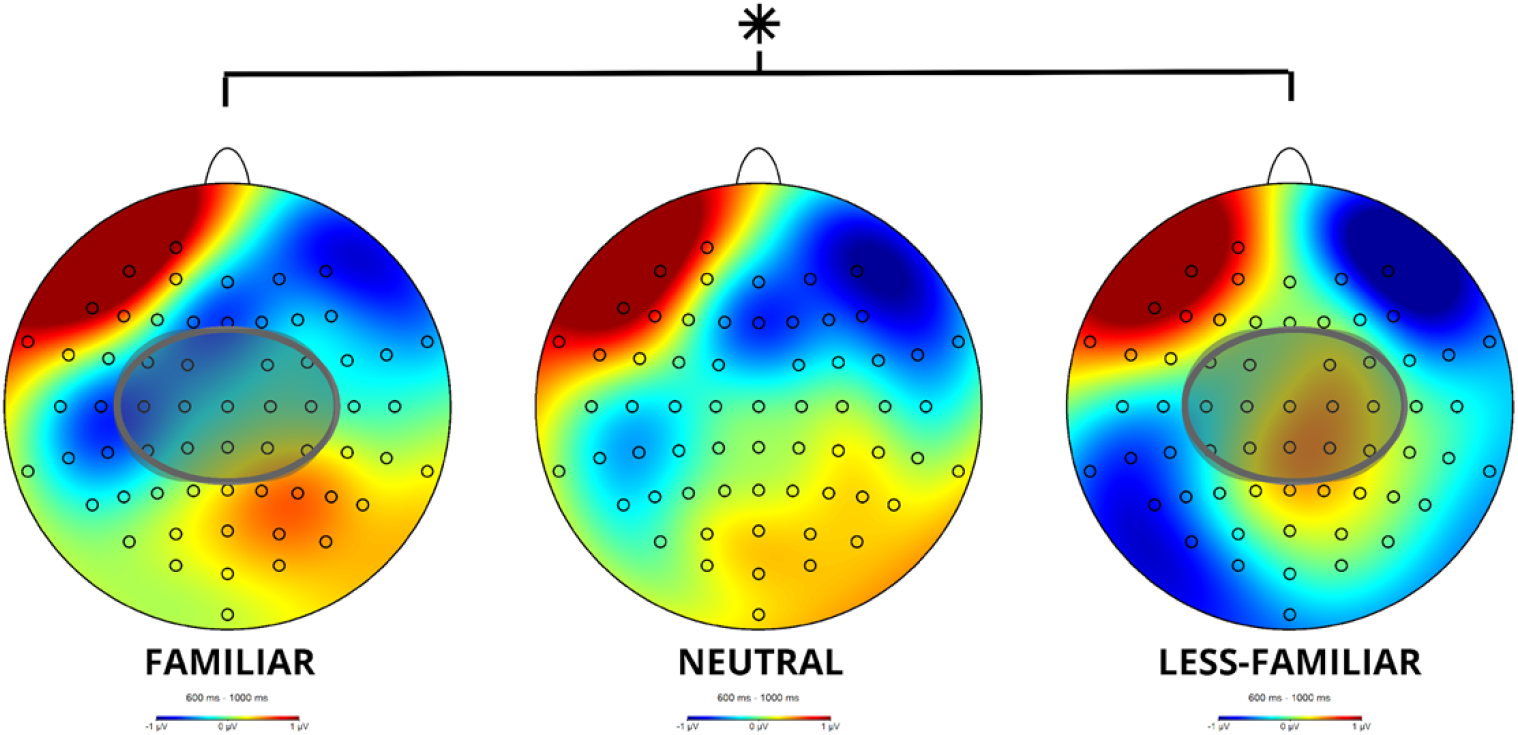
Post N400 component topographical map for locus 2 in the 600-1000 ms time window for the F, N and LF conditions.

**Figure 8.**
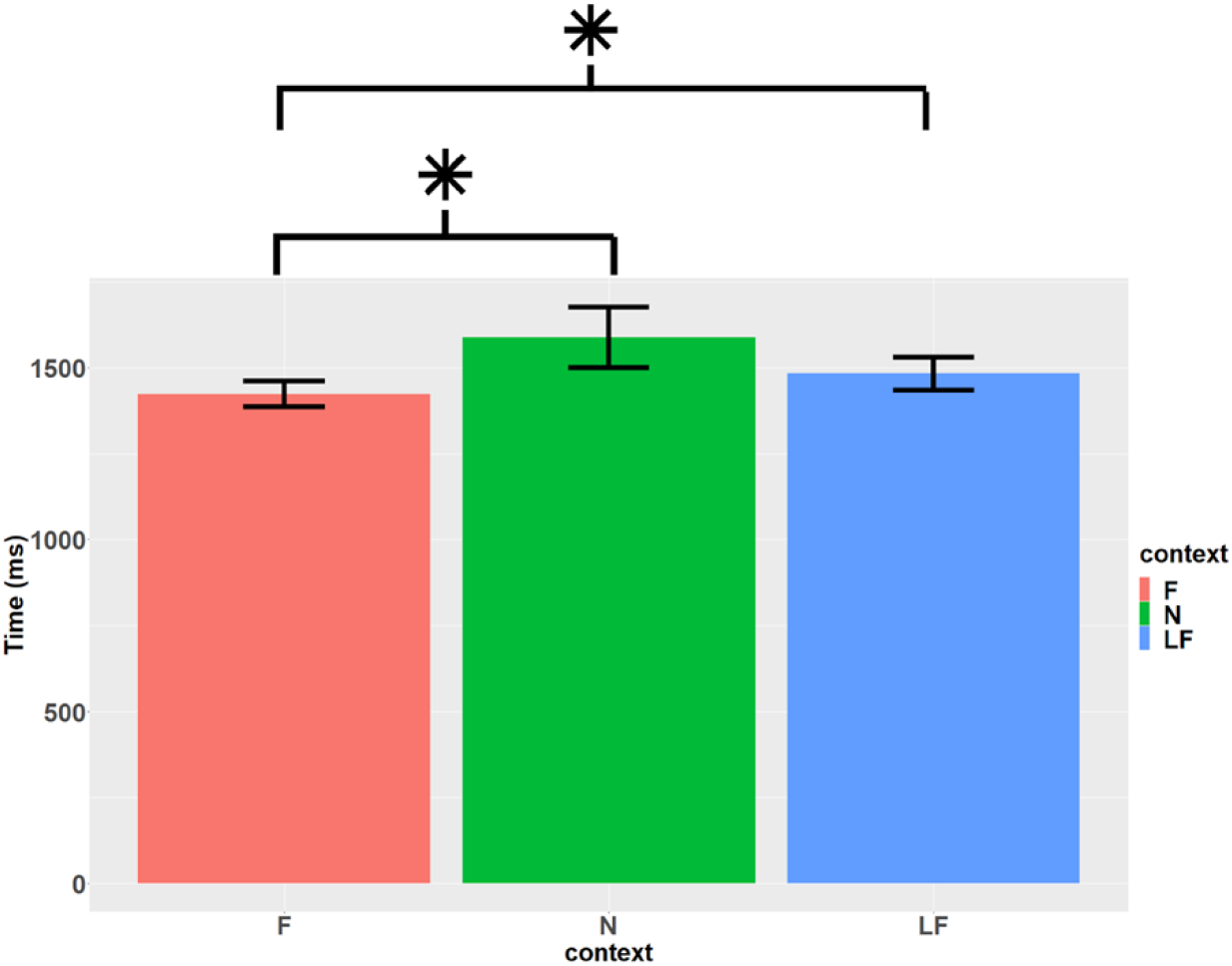
Reaction times according to word context. F(m=1423.7, SD=1069.2, se=36.7), N (m=1588.7, SD=2535.3, se=88.5) and LF (m=1483.4, SD=1382, se=48).

In the case of pseudowords, there is no significant difference associated to context, F(2,2318)=0.56 y p=0.564.

##### Accuracy

When carrying out a likelihood-ratio test, we found that the model that considers context and lexicality predicts accuracy, X^2^(3)=30.2, p<0.001. However, no interaction effect can be found between context and lexicality, X^2^(2)=5.8, p=0.055. Because of this, main effects were analyzed. Additionally, significant effects were found both for context, X^2^(2)=7.65, p=0.02, and lexicality, X^2^(1)=22.68, p<0.001.

When considering context in the Tukey method, a significant difference can be found between the F and N contexts. The odds ratio of responding correctly to F targets is higher (OR=1.70) than to N targets (p=0.02).

Separate lexicality analyses were carried out to look at the differences between words and pseudowords. Regarding context, we found that words are 2.13 times more likely to have a response than pseudowords, p<.001.

Since there is a significant main effect, the lexicality factors were analyzed separately. The context effect on word type showed a significant difference X^2^(2)=13.81,p=0.001. Contrasts reveal a significant difference in the F – N pair, z=-3.44, p=0.00057. This means that the participants were more accurate for F words than for N words, as shown in Figure 9. In the case of pseudowords, there was no significant difference, X^2^(2)=1.82, p=0.4.

**Figure 9.**
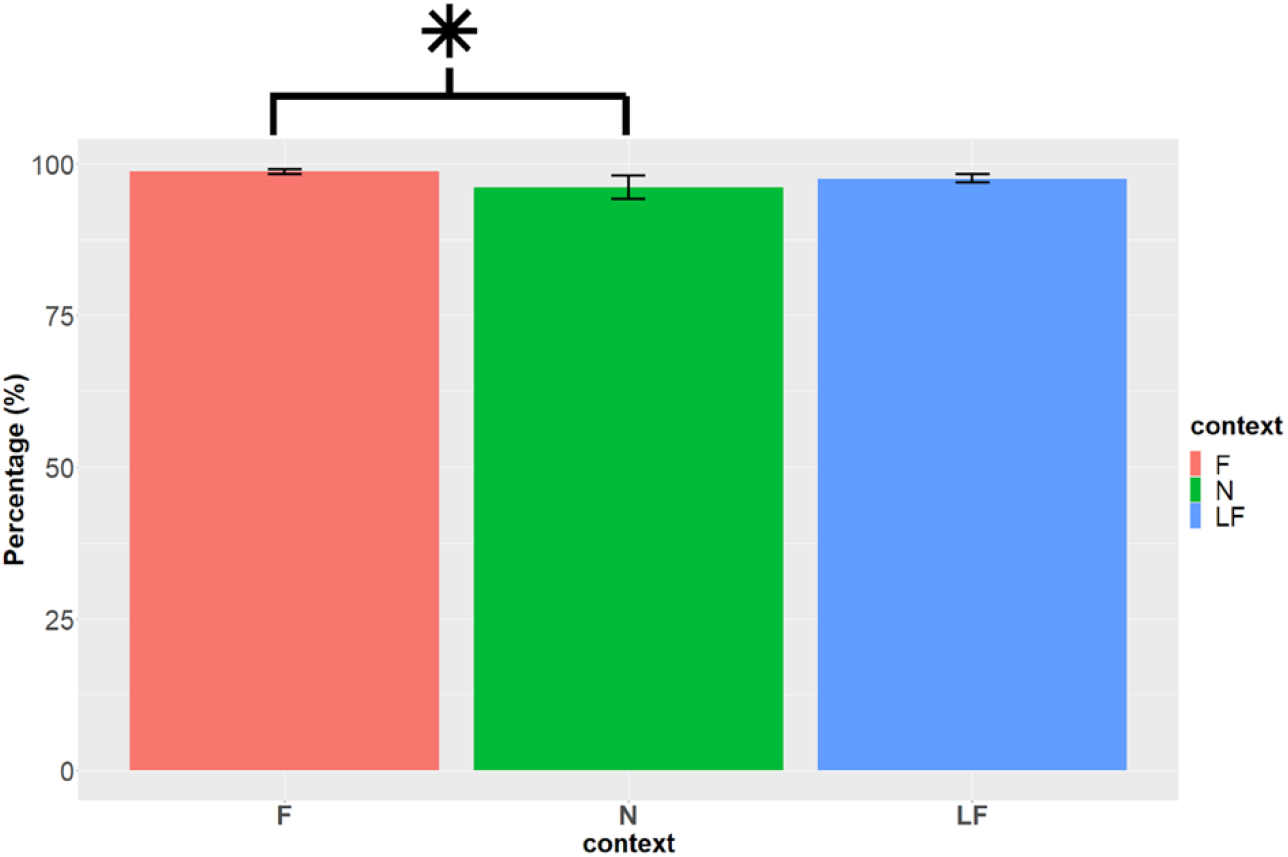
Accuracy (%) according to target word. F (m=98.7, SD=3.2, se=0.42), N (m=96.2, SD=14.6, se=1.93) and LF (m=97.7, SD=5.55, se=0.742).

#### 3.2.1. ERP components: FN400 and N400

##### FN400

For locus 3, an FN400 component in the ROI was found in the 300-500 ms time window, involving electrodes F3, F5, F7, FC3, FC5, FT7. The statistical analysis of the time window showed a significant difference in the context x lexicality interaction, F(2,283.1)=4.05, p=0.019. In order to analyze the effect of context, the lexicality variables were analyzed separately.

There is no significant difference between the means for context (F - N - LF), F(2,113.2)=1.4, p=0.25. When analyzing pseudowords separately, there is a marginal difference, F(2,113.2), p=0.06, between F and LF, t(113)=2.22, p=0.072. The results of the model show a significant difference between F and LF, t=2.22, p=0.029. As shown in *Figure 10* and the topographical map in Figure 12, the amplitude of F context is more positive than that of context LF.

**Figure 10.**
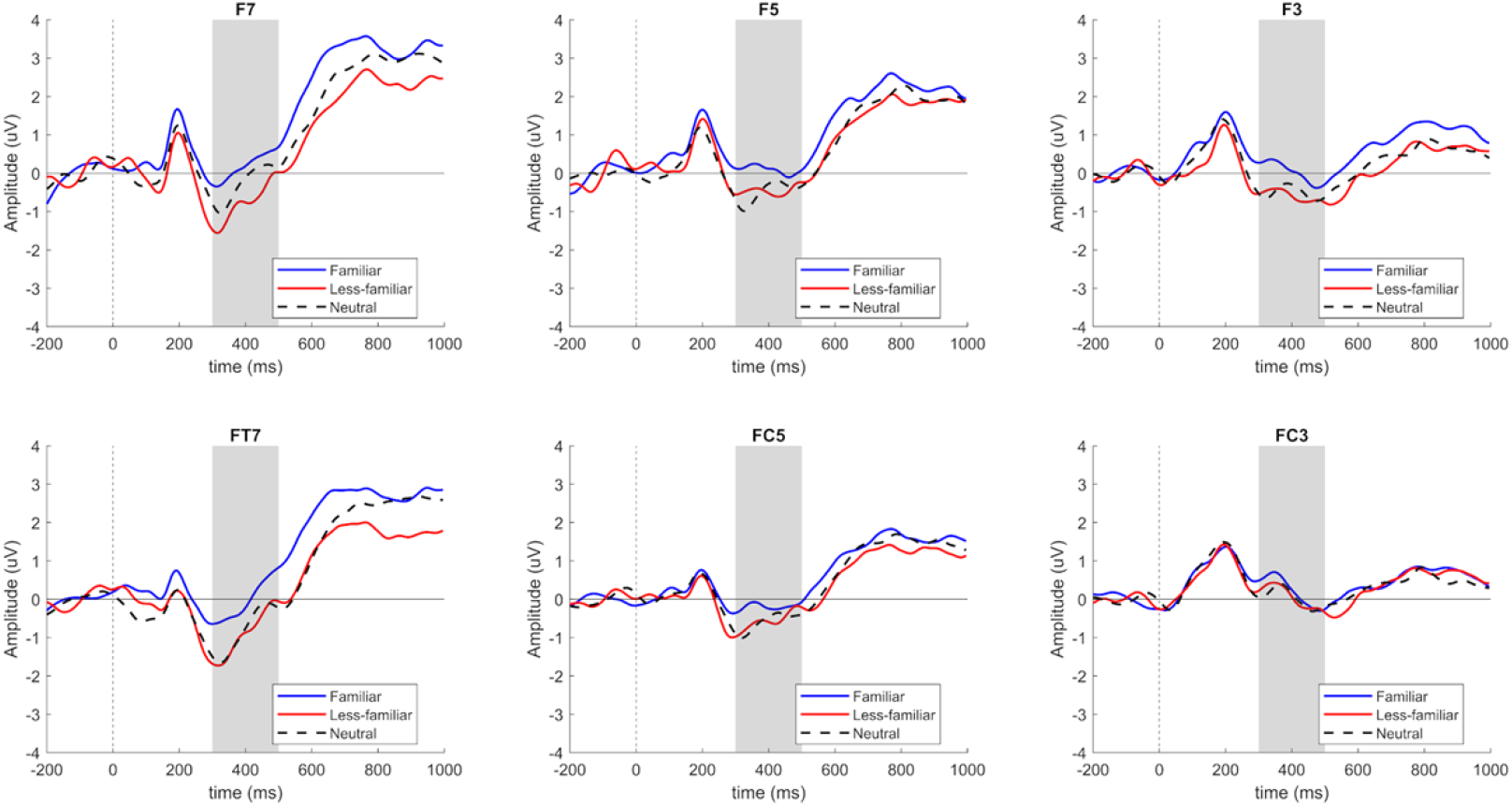
**FN400** component in Locus 3 for pseudowords in electrodes F3 – F5 – F7 – FC3 – FC5 – FT7, sorted by the F, N and LF conditions.

##### N400

In the 300-500 ms time window, there is an N400 component in the ROI involving electrodes C4, C6, CP4, CP6, T8, TP8. Additionally, there is a significant difference in the interaction between the context and lexicality variables, F(2,283.1)=3.64, p=0.027.

When analyzed separately, no significant difference was found for word, F(2,113)=0.32 y p=0.73. On the other hand, the separate analysis for pseudoword showed a significant difference for the context factor, F(2,113)=4.91, p=0.009. Contrasts show a significant difference between F and LF, t(113)=3.11, p=0.002, as shown in *Figure 11* and the topographical map in Figure 12

**Figure 11.**
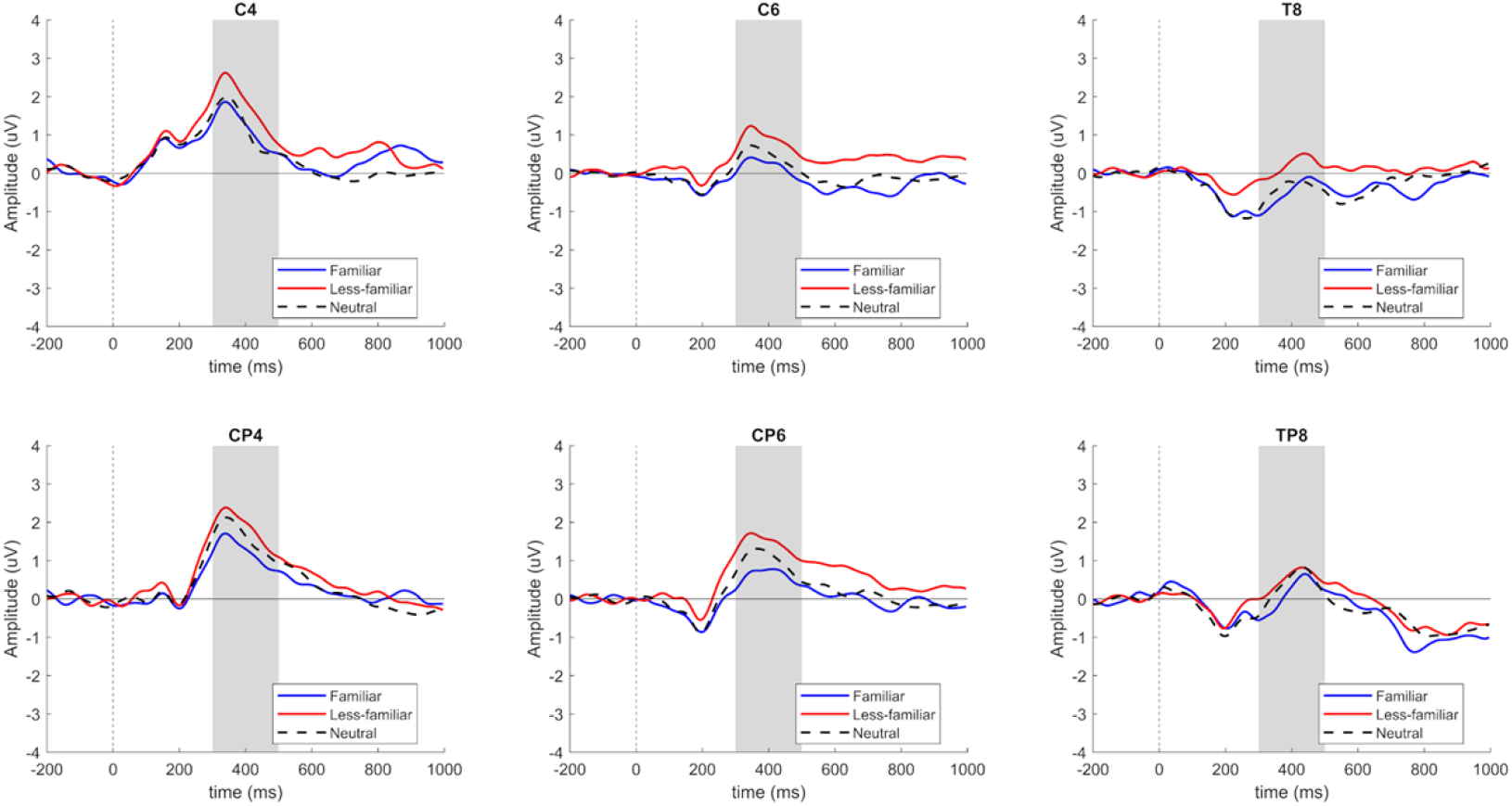
**FN400** component in Locus 3 for pseudowords in electrodes C4 – C6 – CP4 – CP6 – T8 – TP8, sorted by the F, N and LF conditions.

**Figure 12.**
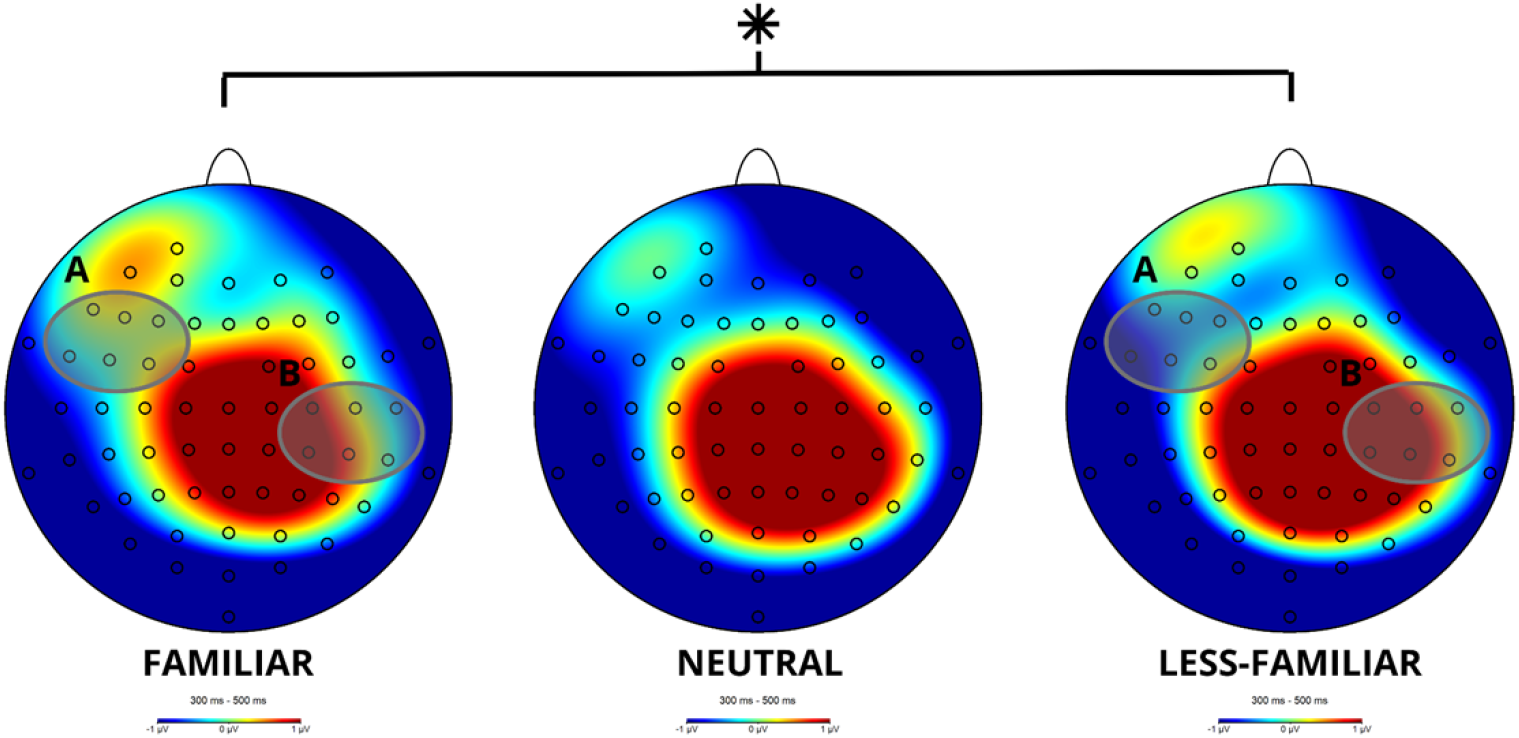
Topographical map of the **FN400** and the **N400** components in Locus 3 for pseudowords in the 300-500 ms time window for the F, N and LF conditions. **A)** ROI involving channels F7, F5, F3, FT7, FC5 and FC3 for the FN400 component; **B)** ROI involving channels C4, C6, T8, CP4, CP6 y TP8 for the N400 component.

## 4. Discussion

The aim of this study was to explore the costs and benefits of the time course of inferences in university students with reading comprehension difficulties. Ours results show the neural time course of inferences in 3 different locus in narration. Firstly, a restricted context, marked by familiarity, a less-familiar context and a neutral context. Secondly, a critical phrase that closes the event proposed within the context. And, thirdly, a target word that is inferred from the story being read.

Our hypothesis posed that the context of the stories would influence students from the start of the narration, causing differences between the familiar and less-familiar contexts via an N400 component. Likewise, we expected to see an influence of lexicality in inference generation, leading to a possible FN400 or N400 effect. Below, we discuss each locus based on which neural component was found and its description in literature.

### LOCUS 1: Slow Negativity Potential component

As hypothesized, context had an influence on university students from the start of the narration. In this locus, we found an unexpected electrophysiological component, the slow negativity potential component. While its topographical distribution is mostly frontal central in some research [45, 46, 47], it has also been found in right dorsolateral regions [48], as was the case in our study. In line with the proposal of some authors [49, 50, 51], the semantic context had an immediate influence on and marks the course of reading.

Some authors [48, 49] have found this potential in two modalities, verbal and auditory, showing the same trend as in our case: higher negativity in a more restricted context, such as highly familiar contexts, when compared to other contexts. In this study, we found statistically significant contrasts between the familiar and less-familiar contexts, as well as between the familiar and neutral contexts, showing larger negativity in familiar contexts. Apparently, this component is associated to the predictability of upcoming events in stories and can be considered an indicator of semantic prediction in top-down discursive processes [50]. It is interesting to note that for the participants of this study, familiar contexts already have a higher prediction cost than other linguistic contexts.

### LOCUS 2: N400 Component

An N400 component was found in the typical time window for that neural correlate, between 300-500 ms, although it had a left and not a right topographical distribution. Reports of this component in literature describe it in both hemispheres with an unstable topographical distribution, as its location depends highly on the experimental manipulation [52, 53]. This left laterality was also reported by Ding et al. [54] for emotion verbs in contrast to neutral words and by Holcomb et al. [55] in unrelated target images. Likewise, Tabullo et al. [16] found a left lateral N400 in a narrower time window: 300-400 ms. The authors manipulated the context (strong/weak) and predictability (expected/unexpected) in two university reader groups: skilled readers and readers with reading difficulties. Group inclusion was based on a Cloze test. The results of the study showed higher negativity for unexpected endings, especially when dealing with restricted contexts, in skilled readers. Additionally, they found a higher positivity associated to the Post-N400 component in unexpected endings, with a higher prominence in the restricted context, only in skilled readers. These results suggest that readers whose reading skill is high made early predictions, based on a more restricted context that manifested in a reduction of the N400 component for strong contexts with a high predictability. Conversely, readers with reading comprehension difficulties showed less effective prediction mechanisms by not manifesting a reduction of the N400 component and by not eliciting late positivity.

In our experiment, it was surprising to find a higher negativity in the familiar critical phrase, instead of in the less-familiar. Apparently, a familiar context elicited a higher cognitive cost, reflected in the N400 component in the moment that the sentence is being processed. It is important to consider that, unlike in Tabullo et al.’s [16] experiment, semantic processing was being registered online, as the inference process has not taken place yet, only the effect of the context on the critical phrase. This phrase reveals a higher negativity in familiar contexts as opposed to a less-familiar context. This result might be due to two plausible reasons: an interference cognitive process that takes place in more concrete sentences, represented in a familiarity context, or difficulties to benefit from the context in order to comprehend the story by less skilled readers. We will address the evidence for both of these reasons.

In the case of concreteness of the phrase, Holcomb et al. [56] found that concrete final words elicited a higher N400 in the 300-500 ms time window and its negativity extended to a later time window, between 500-800 ms. The N400 effect in our study could be explained by the fact that familiar words have a larger amount of semantic information than abstract words. Thus, they have a higher cognitive cost. This effect also takes place in the final locus of counterfactual stories with instrumental type events with a double meaning, which implies the use of a specific tool to achieve a goal [9]. In spite of this, the effect of concrete words seems to dissipate in the context with higher expectations [56], something that does not necessarily happen in our study for the familiar context.

According to Amoruso et al. [57], expectations depend on online contextual information and the reader’s prior knowledge, whose discrepancies are reduced if both constructs coincide. However, a series of experimental studies [58, 59, 60] have reported that in the case of language action, such as what could be found in familiar stories as opposed to less-familiar ones, there is an interference effect when sensorimotor processes take place in parallel to the action. In this case, the familiar stories have a higher sensorimotor representation when compared to less-familiar stories, according to the experience of the reader. This may have caused a higher cognitive cost similar to the interference effect seen in behavioral studies.

Nonetheless, there is a series of studies reporting how readers benefit from context in ERP studies, as previously mentioned in this article [18, 19, 20, 21]. Federmeier et al. [61] found a graduation of context in 3 types of contexts: more restrictive, mildly restrictive and unexpected endings. Their results show an N400 component with a higher negativity in unexpected endings, as opposed to a reduction of the N400 component in more restrictive contexts, when the ending of the sentence adjusted better to the context. Other authors [62] go even further by showing a higher graduation of the contextual level for the final word, in a measure that proposed five ranges of likelihood, ranging from 10 to 30%, 30 to 50%, 50 to 75%, 75 to 90% and 90 to 100%, as opposed to an unexpected ending. Results showed an N400 effect with higher negativity in the unexpected range and a graduation of such negativity according to the five ranges of likelihood, according to the reader’s expectations. Thus, the amplitudes of the N400 component decreased inversely to the increase of restriction.

Previous studies show this context graduation taking the opposite direction to that of our study. Because of this, it is worth questioning whether predictive processing associated to context takes place under every circumstance or whether the processes involved in the inhibition and/or review of a prediction in more familiar contexts might be failing in readers with reading comprehension difficulties. It is likely that our students’ processing is rather bottom-up, based on the integration of lexical and semantic information, rather than interactive, where context information is used in advance or “predictively”. This second interpretation of the results will be explained below.

There are few studies that delve into the individual differences in reading comprehension [16]. Individual differences have been found in bilingual groups with higher or lower efficiency, where the latter group showed a greater amplitude of the N400 component and a higher late positivity (LPC component) in sentences with an explicit continuation or paraphrasing, instead of more conceptual variables like inferences [63]. Another study [64] showed differences between participants with high working memory and those with lower working memory. The results showed that only the former group was able to correct an incorrect contextual interpretation and integrate new information through a higher negativity of the N400, while lower working memory participants faced difficulties to inhibit the initial interpretation and to integrate it into the discourse through a lesser amplitude of the N400 component.

A more specific study, Landi and Perfetti’s [65], showed how skilled readers benefit from the context in a verbal semantic pairing task, where participants had to choose whether a pair of words were related or not. The results showed a higher negativity in the N400 component for semantically unrelated pairs in both populations. However, the less-skilled readers showed a reduction of N400. Additionally, the polarity of the N400 in this task showed higher negativity in less-skilled readers when compared to skilled readers. This pattern was repeated in pairing tasks involving word-picture pairs and in a phonological task, i.e., deciding whether two words are pronounced in the same manner. This negativity has not been explored in depth. However, it can be seen in our study for the N400 component and the post-N400 component, where polarity associated to the experimental conditions remains more negative in the experimental condition associated to familiarity, displaying the opposite direction to what would be expected. In other words, there is a higher positivity for less-familiar contexts when compared to familiar ones.

The post-N400 component is a reflection of the cost of prediction error that takes places due to a higher inhibition of the preceding words [66, 67]. This component arises when there is a conflict between the reader’s expectations and the construction of the situation model in more restrained contexts [16, 49, 61, 68, 69]. However, the higher positivity in our experiment was found in less-familiar contexts. In this position, we expected to find a P600 component, similarly to the one found in Steele et al.’s [7]. Nevertheless, the P600 component takes place at an earlier stage, in the 400-800 ms time window, showing a higher deflection at around 600 ms. The post-N400 component has a larger distribution, including frontal, central and parietal regions, which reflect additional processes of semantic integration, such as the resolution of semantic conflicts or re-analyses related to lexical anticipation processes [66]. Such cognitive processes seem to be derived from the processing of less-familiar contexts as opposed to familiar ones.

### LOCUS 3: Lexical decision task

In the third locus of this study, we can truly see when textual inference takes place, since in a lexical decision task readers decide if the following lexical entry is a word or a pseudoword. The word was derived from both contexts, familiar and less-familiar, and did not hold any relation to the neutral context. Pseudowords shared the same phonological structure with the words, as only one syllable was changed and the consonants were replaced by others similar to the ones in the target. Behavioral results of this research reveal no significant interaction effect in reaction times or accuracy, although there was an effect of lexicality leaning toward longer times and lesser accuracy for pseudowords.

By analyzing words separately, just as in Steel et al.’s [7] study, wherein pseudowords were excluded from the analysis, we found significant differences between the conditions. These differences were evidenced by shorter reaction times for familiar words than for less-familiar and neutral words. Likewise, we found higher accuracy rates for familiar words when compared to neutral ones. These results show that readers benefited from context in order to make an inference. It stands out that in Steele et al.’s study [7], words derived from the less-familiar context took less time than words derived from the familiar context, although the difference in contrasts took place between the familiar and the neutral context and between the less-familiar and the neutral context. In our study, we did not find any significant differences between the familiar and the less-familiar context. On the other hand, it is interesting that reaction times of our participants were considerably longer to that of Steele et al.’s [7] participants. In their results, the authors reported mean reaction times of 570 ms for the familiar condition, as opposed to 1423 ms for the familiar condition reported in our experiment. This was also the case for the other variables in which less-skilled readers virtually took twice as long.

In the electrophysiological analysis, we found a statistically significant interaction in the FN400 component. However, in the separate analysis, we only found significant differences in the familiar and less-familiar contrasts for pseudowords. As mentioned above, the FN400 component is a neural correlate of familiarity [22, 23, 24, 25, 26]. Due to this, we expected to find some effect on this component. Nevertheless, lexicality seems to interact with context and less-skilled readers paid special attention to pseudowords that kept the same syllabic structure of slightly modified words. In this regard, we found an attenuation of the FN400 component in the familiar context, as opposed to less-familiar contexts, as is usually reported in literature for the case of words. Facilitation by context of pseudowords can be explained by Perfetti’s [70, 71] theory, wherein lexical access in less-skilled readers is very slow, as reflected by their reaction times. This causes said readers to have enough time for contextual information to start to flow on lexical access.

In regards to the N400 component, statistically significant effects were once again found for the context x lexicality interaction, although effects occur only for pseudowords. In our case, there is greater negativity in the less familiar context when compared to the less-familiar, while in Steele et al.’s [7] case, there was a higher negativity in neutral contexts when compared to the familiar and less-familiar experimental conditions for words. Finally, the authors found the P600 component in this position, revealing a reinterpretation effect for familiar and less-familiar contexts in connection with neutral contexts, but in our case there was no late effect in the third locus.

An important difference between Steel et al.’s [7] study and our study is in the SOA interval between the final phrase and the target word, since the researchers were looking for predictive inferences in a 1000 ms inter-stimulus timeframe. Conversely, in our study, we were looking for automatic inferences, without inter-stimulus interval, in order to record bridging or forward inferences establishing causal coherence. Moreover, it is worth noting that the sample of this study was made up of less-skilled readers, who have difficulties to make inferences connecting distant contents in a text [72].

On the other hand, the restrictions of the familiar context in a self-administered task, due to the process of reading and the presentation time of the target word, favor the generation of online inferences [73, 74, 74]. Support for this notion can be found in McKoon and Ratcliff [31, 32], who posit that such inferences are generated if information is readily available from general knowledge, as is the case in familiar contexts. The familiarity effects occurs for words in the behavioral task and is different from the original Study [7], where the less-familiar context saw higher facilitation in reaction times for the target word, which took place within time limits, as opposed to our results.

Further evidence that inferences were generated online is that pseudowords behaved virtually like real words as per the FN400 component. According to Rodríguez [64], the ability to automatically make inferences extends to pseudowords and can affect an adequate inference on words, which are more semantically grounded on discourse. The N400 component shows the cognitive cost of familiarity by eliciting higher negativity in less-familiar pseudowords when compared to familiar ones, regardless of the prior context in which familiarity could have attenuated the N400 component.

However, it is important to note that readers have reading comprehension difficulties. This is reflected by the absence of the P600 component, which was not observed in Steele et al.’s [7] experiment. The absence or attenuation of this component has also been found in other research on less-skilled readers [16, 63, 67]. The lack of this component in our study, in addition to the findings in locus 1 through locus 3, seem to account for a predominantly bottom-up processing instead of a top-down processing in less-skilled readers. Apparently, lexical effects modulate cognitive processing, both in locus 2 and in locus 3. In locus 1, however, readers seem to benefit from context.

According to current theories on discourse comprehension, reading processes are partially incremental, flexible and dependent on context [52]. For this reason, there is a high likelihood that, based on the context declared in locus 1 of this research, some semantic content may have generated an expectation from a word. In turn, this may have generated a conflict in the following locus through the N400 and post-N400 components [76]. As a result, a combination of both types of processing—top-down in locus 1 and locus 3 in the behavioral task, and bottom-up in locus 2 and 3 in electrophysiology— may coexist in the case of less-skilled readers, who do not control which strategy is the most effective in each locus. Good readers, on the other hand, use preferentially top-down processing, benefiting from semantic networks in the use of context, as well as bottom-up processing [16].

## 5. Conclusions

Ultimately, the time course of inferences in less-skilled readers shows the benefits of context through a use of a slow negativity potential component in the first locus, which is associated with higher negativity in familiar contexts. Conversely, in the second locus we see the cognitive cost that familiar context entails, eliciting a higher negativity in the N400 and post-N400 component. The latter component has been associated with lexical anticipation processes rather than contextual anticipation processes of discourse [66]. Lastly, in the third locus, participants benefited from context in the behavioral task by generating automatic inferences associated with a higher facilitation of words coming from a familiar context, as opposed to a less-familiar or neutral one. Nonetheless, effects can be seen in electrophysiological components, fundamentally in pseudowords, showing the benefit of context for such lexical entries, as well as the cost of lexicality in the semantic processing of words.

One of the limitations of this study is not having a group of skilled readers to establish a direct contrast between both populations. This implies that all interpretations of this study must be taken with care, as no comparisons can be made with the current sample. An additional limitation is the lack of SOA graduation studies, i.e. two experiments with the same population comparing a short SOA with a long SOA in order to establish a clearer difference between automatic and predictive inferences. In future research, the combination of electrophysiological techniques —used mainly in contextual semantic processing— with eye movement technique, allowing for a better monitoring of lexical and even sublexical processes, might shed more light on the costs and benefits of experimenting with less-skilled readers, as opposed to skilled readers.

## Author Contributions

Conceptualization, Urrutia, M..; methodology, Urrutia, M., Pino, E..; software, Pino, E., Troncoso-Seguel.; formal analysis, Troncoso-Seguel, M., Bustos, C..; investigation, Torres-Ocares, K., Mariángel, S.; resources, Urrutia, M., Pino, E., Guevara, P..; data curation, Bustos, C..; writing—original draft preparation, Urrutia, M.; writing—review and editing, Marrero, H. Yang, F..; visualization, Troncoso-Seguel, M., Torres-Ocares, K..; supervision, Urrutia, M., Pino, E., Guevara, P.; project administration, Urrutia, M.., Pino, E.; funding acquisition, Urrutia, M.. All authors have read and agreed to the published version of the manuscript.

## Institutional Review Board Statement

The study was conducted in accordance with the Declaration of Helsinki, and approved by Ethics, Bioethics and Biosafety Committee of Universidad de Concepción (Chile) (Protocol No. CEBB906-2021) for studies involving humans.

## Informed Consent Statement

Informed consent was obtained from all subjects involved in the study.

## Data Availability Statement

data is unavailable due to privacy or ethical restrictions.

## Acknowledgments

Thanks to María Fernanda Cornejo Rodríguez and Ana Belén Parra Alarcón for their contributions to data collection.

